# Zoledronate extends healthspan and survival via the mevalonate pathway in a FOXO-dependent manner

**DOI:** 10.1101/2020.04.09.033498

**Authors:** Zhengqi. Chen, Julia Cordero, Cathy Slack, Martin Zeidler, Ilaria Bellantuono

**Affiliations:** Healthy Lifespan Institute, Department of Oncology and Metabolism, The Medical School, Beech Hill Road, University of Sheffield, Sheffield S10 2RX; Institute of Cancer Sciences, University of Glasgow, Beatson Institute for Cancer, Research Switchback Rd, Bearsden, Glasgow G61 1QH; School of Life & Health Sciences, Aston University, Aston Triangle, Birmingham B4 7ET; Department of Biomedical Science, The University of Sheffield, Western Bank, Sheffield S10 2TN

**Keywords:** aging, FOXO, healthspan, lifespan, bisphosphonates, DNA damage, *Drosophila*

## Abstract

Increased longevity has not been paralleled by extended healthspan, resulting in more years spent with multiple diseases in older age. As such, interventions to improve healthspan are urgently required. Zoledronate is a nitrogen containing bisphosphonate, which inhibits the farnesyl pyrophosphate synthase (FPPS) enzyme, central to the mevalonate pathway. It is already used clinically to prevent fractures in osteoporotic patients, who have been reported to derive unexpected and unexplained survival benefits. In this study we show that zoledronate has beneficial effects on both lifespan and healthspan using *Drosophila* as a model. We found that zoledronate extended lifespan, improved climbing activity and reduced intestinal epithelial dysplasia and permeability in aged flies. Mechanistic studies showed that zoledronate conferred resistance to oxidative stress and reduced accumulation of X-ray-induced DNA damage via inhibition of FPPS. Moreover, zoledronate inhibited pAKT in the mTOR pathway and functioned via dFOXO, a molecule associated with increased longevity, downstream of the mevalonate pathway. Taken together, our work indicates that zoledronate, a drug already widely used and dosed only once a year to prevent osteoporosis, modulates important mechanisms of ageing. Its repurposing holds great promise as a treatment to improve healthspan.

## INTRODUCTION

It has been estimated that by 2050, Europe, North America, and eight countries across the other continents will have more than 30% of their population over the age of 60 (World Health Organization, 2015). However, this increase in life expectancy has not translated into extended healthspan (Bellantuono, 2018). More than 60% of those over the age of 65 suffer from multimorbidities – defined as the co-existence of multiple chronic health conditions (Hung, Ross, Boockvar, & Siu, 2011; Marengoni et al., 2011; Vogeli et al., 2007). These multimorbidities impose quality-of-life and care management challenges associated with treating multiple conditions individually and are frequently associated with increased costs, reduced efficacy and increased likelihood of adverse events due to polypharmacy (Gandhi et al., 2003; JH, TS, LR, & al, 2003; Tinetti, Bogardus, & Agostini, 2004). Targeting pathways underpinning central mechanisms of ageing which are common to multiple disorders offers new opportunities to treat multimorbidity and overcome these problems.

Zoledronate (Zol) is a nitrogen containing-bisphosphonate used for the treatment of skeletal disorders, including osteoporosis. It inhibits the mevalonate pathway through inhibition of the enzyme farnesyl pyrophosphate synthase (FPPS) (Amin et al., 1992; Luckman, Hughes, Coxon, Russell, & Rogers, 1998) and thereby inhibits bone resorption by osteoclasts. Due to its high affinity binding to hydroxyapatite crystal mineralisation, Zol is stored in bone and released following bone breakdown conferring long lasting activity (Weiss et al., 2008). In clinical practice, Zol is administered by intravenous infusion once a year in post-menopausal women to prevent fractures (Lambrinoudaki, Vlachou, Galapi, Papadimitriou, & Papadias, 2008; McClung et al., 2007). Recently, retrospective analysis of several clinical trials showed that patients treated with Zol had a significant decrease in mortality rates (Bolland, Grey, Gamble, & Reid, 2010; Colon-Emeric et al., 2010; Eriksen et al., 2009; P. Lee et al., 2016b; Lyles et al., 2007; Morgan et al., 2010). In addition, among patients admitted to intensive care units, those pre-treated with Zol had increase survival despite overall higher level of multimorbidities and older age (P. Lee et al., 2016b). It is unknown whether other mechanisms independent from bone protection may be involved in the Zol-dependent extension of survival.

The mevalonate pathway is an important metabolic pathway responsible for the production of cholesterol and protein prenylation. One key group of isoprenylated proteins are small GTPases (Zhang & Casey, 1996), important signalling molecules, which play central roles in multiple cellular processes, including cellular morphology, integrin function and longevity-associated pathways such as mTOR (Fukata, Nakagawa, & Kaibuchi, 2003; Lawson & Burridge, 2014; Nguyen, Frank, & Jewell, 2017; Park & Bi, 2007). In this study we examined whether Zol was able to extend lifespan and healthspan independent of its effect on bone and whether it was doing so through mechanisms related to ageing, using *Drosophila*, a model widely accepted for ageing studies. While retaining evolutionarily conserved components of both the mevalonate and mTOR pathways, it does not feature the bone-like mineralisation present in mammalian models (Moussian, Schwarz, Bartoszewski, & Nüsslein-Volhard, 2005), so allowing osteo-protective and gero-protective activities to be dissected *in vivo*. Our results uncovered a previously unrecognised role of Zol extending both lifespan and healthspan through dFOXO signalling, a highly conserved molecule well known for its positive effects on longevity.

## RESULTS

### Zol increases lifespan in *Drosophila*

To determine whether Zol might have beneficial effects on the lifespan of *Drosophila*, both male and female *w^Dah^* flies were maintained on standard fly food supplemented with 1μM or 10μM Zol from 4-day of adult life onwards (Fig 1A&B). The survival of males fed with 1μM Zol was significantly increased compared to the vehicle-treated group (Fig. 1A). By contrast, females, showed no significant beneficial effect when treated with Zol at 1μM throughout their lives (Fig. 1B) while 10μM of Zol had adverse effects on female survival (Fig. 1A&B).

**Fig. 1.**
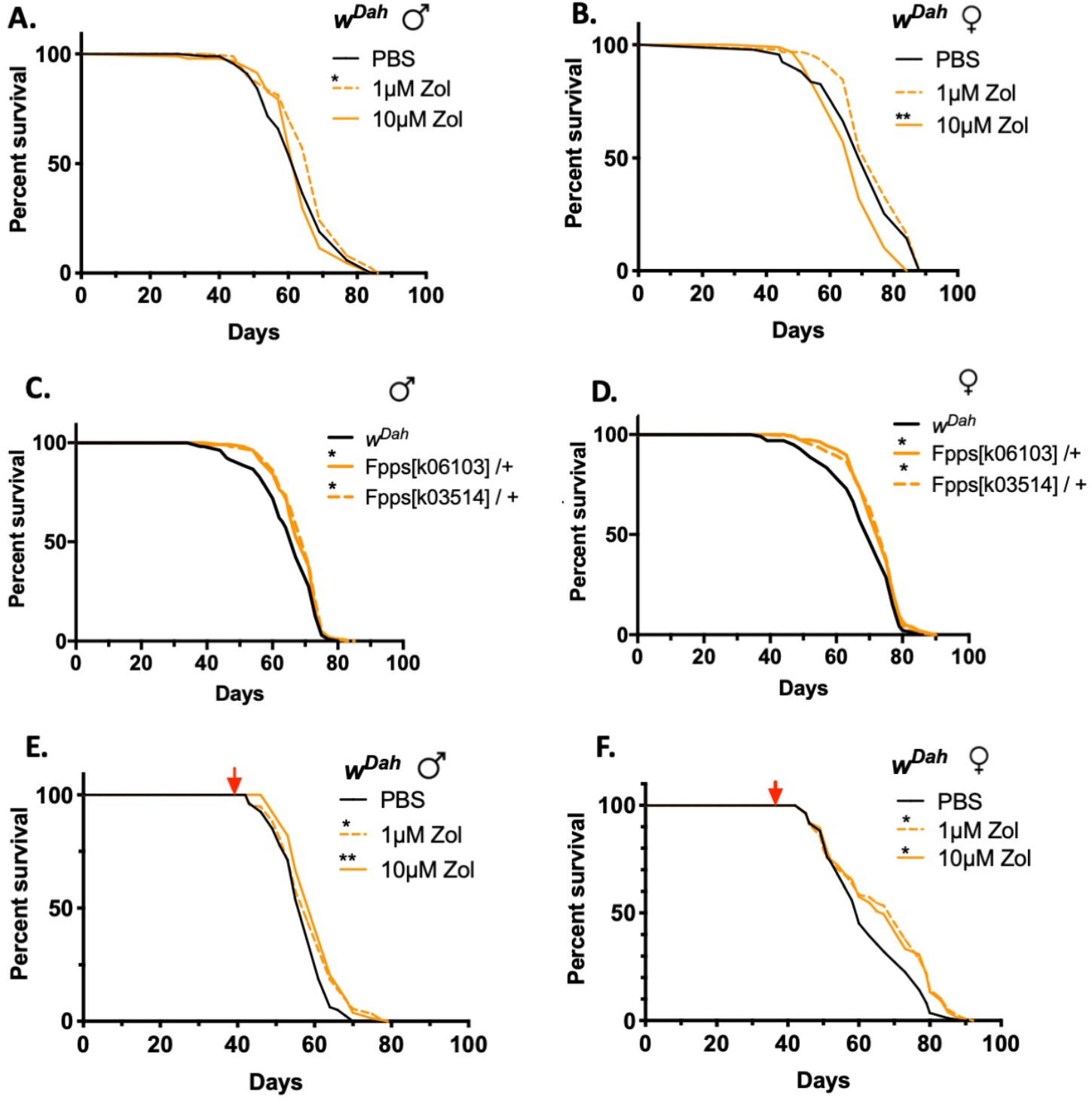
Administration of Zol affects lifespan of flies. Percentage survival of (A) male and (B) female *w*^*Dah*^ flies fed with food in presence or absence of Zol (1 or 10μM) throughout their lives. Percentage survival of (C) male and (D) female heterozygote FPPS mutant flies and *w*^*Dah*^ fed with standard Drosophila food. Percentage survival of (G) male and (F) female *w*^*Dah*^ flies fed with food in presence or absence of Zol (1 or 10μM) from 40 days old. Log-rank (Mantel-Cox) test in Graphpad Prism used to statistically analyse the survival curves, * P ≤ 0.05, ** P ≤ 0.01, *** P ≤ 0.001, **** P ≤ 0.0001

As Zol modulates the mevalonate pathway through inhibition of FPPS, we next tested the lifespan of two heterozygous FPPS mutants (*Fpps*^*k06103*^/+ and *Fpps*^*k03514*^/+) – two independently generated homozygous-lethal loss-of-function alleles containing transposon insertions within the FPPS gene (Spradling et al., 1999). Following at least 7 generations of outcrossing into the *w*^*Dah*^ genetic background, both mutant strains showed a significant increase in overall survival in males and females compared to *w*^*Dah*^ controls (Fig. 1 C&D), suggesting that reduced FPPS activity mediates an extension in lifespan

To limit potential side effects due to long term drug treatment, we next assessed the effects of Zol on lifespan when starting administration at day 40 of adult life (middle age) (red arrow in Fig. 1E&F). This late treatment led to a significant increase in survivorship in both males and females for both 1 and 10μM Zol compared to controls (Fig. 1 E&F). This lifespan extension was significant for both males and females with 1μM Zol producing average median lifespans increases of 1.79 % (±0.10%) and 11.53% (±2.10%)(N=3) for male and females respectively, while in 10μM Zol, males increased by 4.67% (±1.03%) and females by 16.51% (±4.94%) (N=3). Finally, to verify that these results were not influenced by calorific restriction caused by unpalatable food we exposed the flies to drug-laced food containing a non-toxic food colorant via which the volume of food consumed can be measured. No significant difference in food uptake was observed with any of the drugs used (Fig. S1 n=60/trial, 2 trials), suggesting that the extension of lifespan observed is a consequence of Zol treatment.

### Zol increases healthspan in *Drosophila*

To determine whether Zol also improves signs of healthspan we next tested its effects in *Drosophila* by assessing their ability to climb and by the presence of signs of intestinal epithelial dysplasia and intestinal permeability. Displaying strong negative geotactic responses, climbing activity is a widely used assay of *Drosophila* health and activity (Gargano, Martin, Bhandari, & Grotewiel, 2005; Haiyan Liu et al., 2015) and was assessed using the rapid iterative negative geotaxis (RING) assay (Fig. 2). We classed as “high climbers” flies able to climb over 10cm in 15 seconds. As expected, the percentage of high climbers significantly decreased with age in both males and females (Fig 2). Strikingly, an increase in climbing ability was observed in both males and females maintained on food containing Zol from day 4 of adult life. Statistically significant increases in climbing are measured in females at day 42 and in males at day 63 (Fig. 2 A-B). For flies treated with Zol from day 40 of adult life, an increase in climbing ability was observed in both males and females but reached statistical significance in males at day 56 and females at both day 56 and 63 (Fig. 2 C-D). However, no beneficial effect of Zol was observed at a later time point in any of the conditions. These data show there is an overall improvement in climbing ability independent of effects on lifespan.

**Fig. 2.**
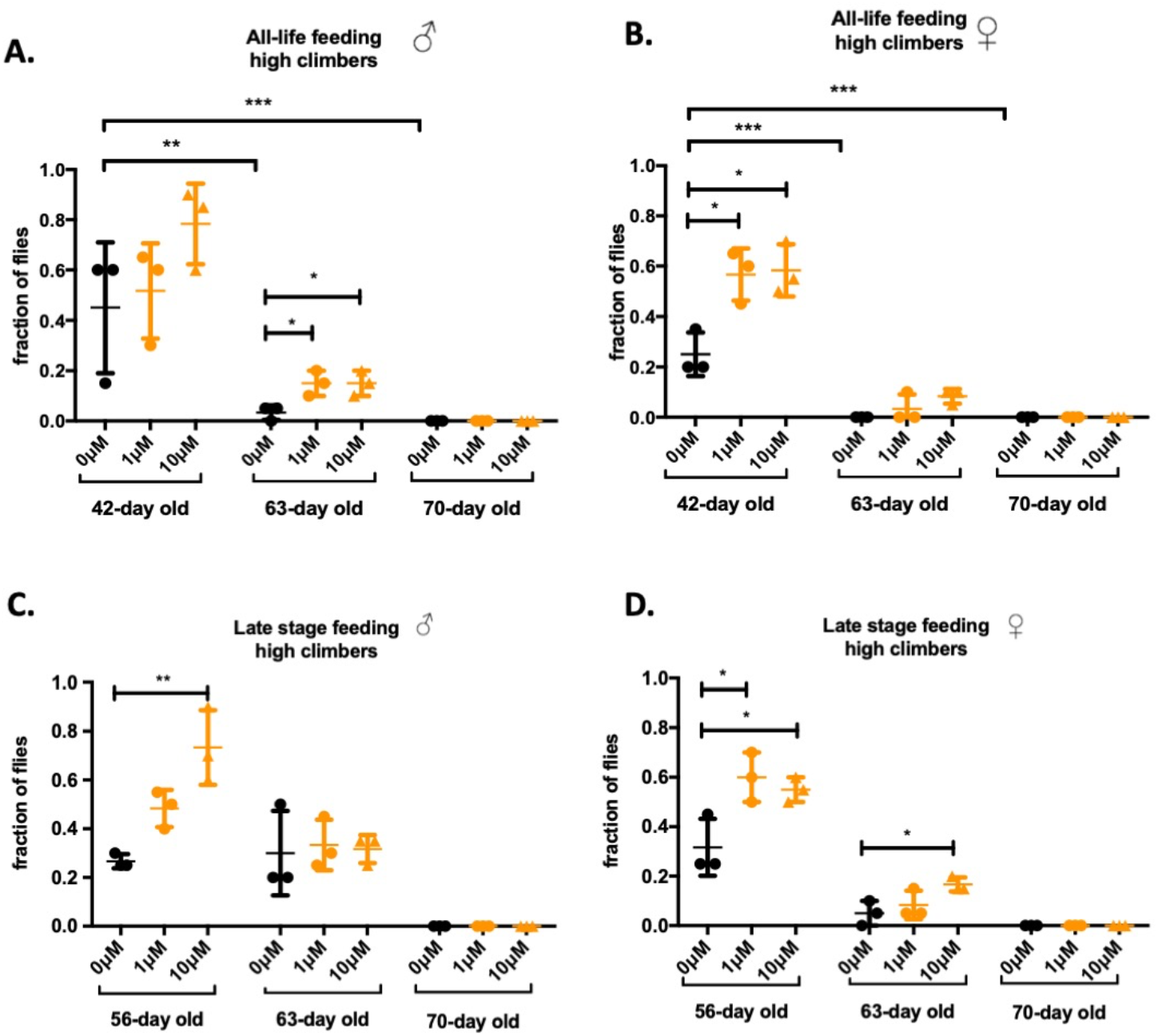
Flies fed with Zol shows a significant increase in climbing ability at later stage of their lives. Lifelong treatment with Zol: (A) Percentage of male flies climbed above 10cm (high climbers) within 15 seconds. (B) Percentage of female flies climbed above 10cm (high climbers) within 15 seconds. Flies treated with Zol from midlife: (C) Percentage of male flies climbed above 10cm (high climbers) within 15 seconds. (D) Percentage of female flies climbed above 10cm (high climbers) within 15 seconds. * P ≤ 0.05, ** P ≤ 0.01, *** P ≤ 0.001.

As flies age, intestinal barrier dysfunction and epithelial dysplasia develop in female flies, a development associated with increased mortality (Martins, McCracken, Simons, Henriques, & Rera, 2018) and considered a good markers of healthspan. To determine the effects of Zol on epithelial dysplasia we examined intestinal morphology at day 7, 42 and 63 of adult life following treatment with vehicle or Zol starting at either day 4 or 40 of adult life. Using *Su(H)-lacZ; esg-Gal4, UAS-GFP* reporters we labeled intestinal stem cells and enteroblasts (progenitor cells) on the basis of GFP expression driven by the *escargot* promoter-a marker associated with stemness. Epithelial dysplasia in the *Drosophila* gut is characterised by hyperproliferation, as identified by the G2/M marker phospho-histone3 (PH3) (Ren et al., 2010) and mis-differentiation of intestinal stem cells (Biteau et al., 2010; Biteau, Hochmuth, & Jasper, 2008). As expected, we observed an increase in intestinal stem-cell proliferation with age in control flies as shown by the significant increase in the number of GFP+ / PH3 positive cells with age (Fig 3A and B) and an increase in the overall proportion of GFP+ cells (Fig.3 C and D). Both parameters are significantly decreased in female flies treated with Zol (Fig.3A-=D).

**Fig. 3.**
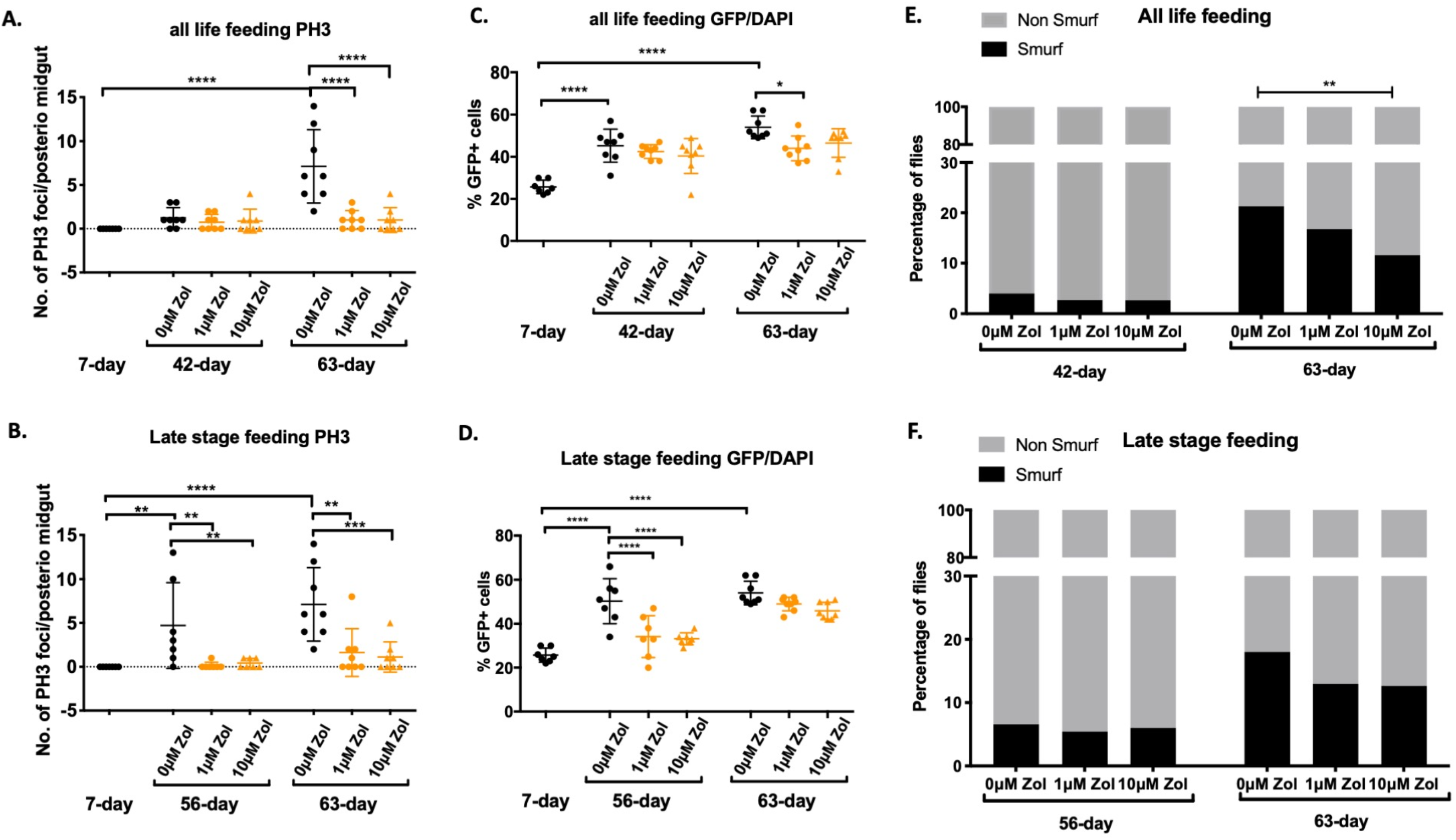
Treatment with Zol reduces epithelial dysplasia and intestinal permeability in *Drosophila*. (A-B) Quantification of PH3+ cell in the intestine of *Drosophila*. Flies were treated with or 1 or without 10μM of Zol from 4 days of age (A) or 40 days of age (B). At least 7 midguts were assessed per condition and per time point. (C-D) Quantification of %GFP+ cells (normalised to DAPI) in the intestine of flies treated with or without 1 or 10μM of Zol from 4 days of age (C) or 40 days of age (D). (E-F) Percentage of flies Smurf positive following treatment with or without 1 or 10μM of Zol starting from 4 days of age (E) or 40 days of age (F) (n>65 per condition per timepoint). Data were analysed with one way ANOVA and Bonferroni post-hoc test *p<0.05, **p<0.01, ***p<0.001, ****p<0.0001

One of the key roles played by the intestinal epithelia is to provide an impermeable barrier to the exterior environment in which damaged or dying epithelial cells are replaced with new cells generated by the ISCs. A loss of barrier function in the intestinal epithelium has been reported with age in both flies and humans. To assess intestinal integrity we performed the “Smurf assay” (Rera, Clark, & Walker, 2012) in which blue colouring added to regular food is able to cross a compromised epithelium and stain the entire fly blue. Consistent with previous findings, the percentage of flies presenting the ‘Smurf’ phenotype was increased with age and is attenuated by treatment with Zol under the all life feeding regime (Fig.4E-F).

**Fig. 4.**
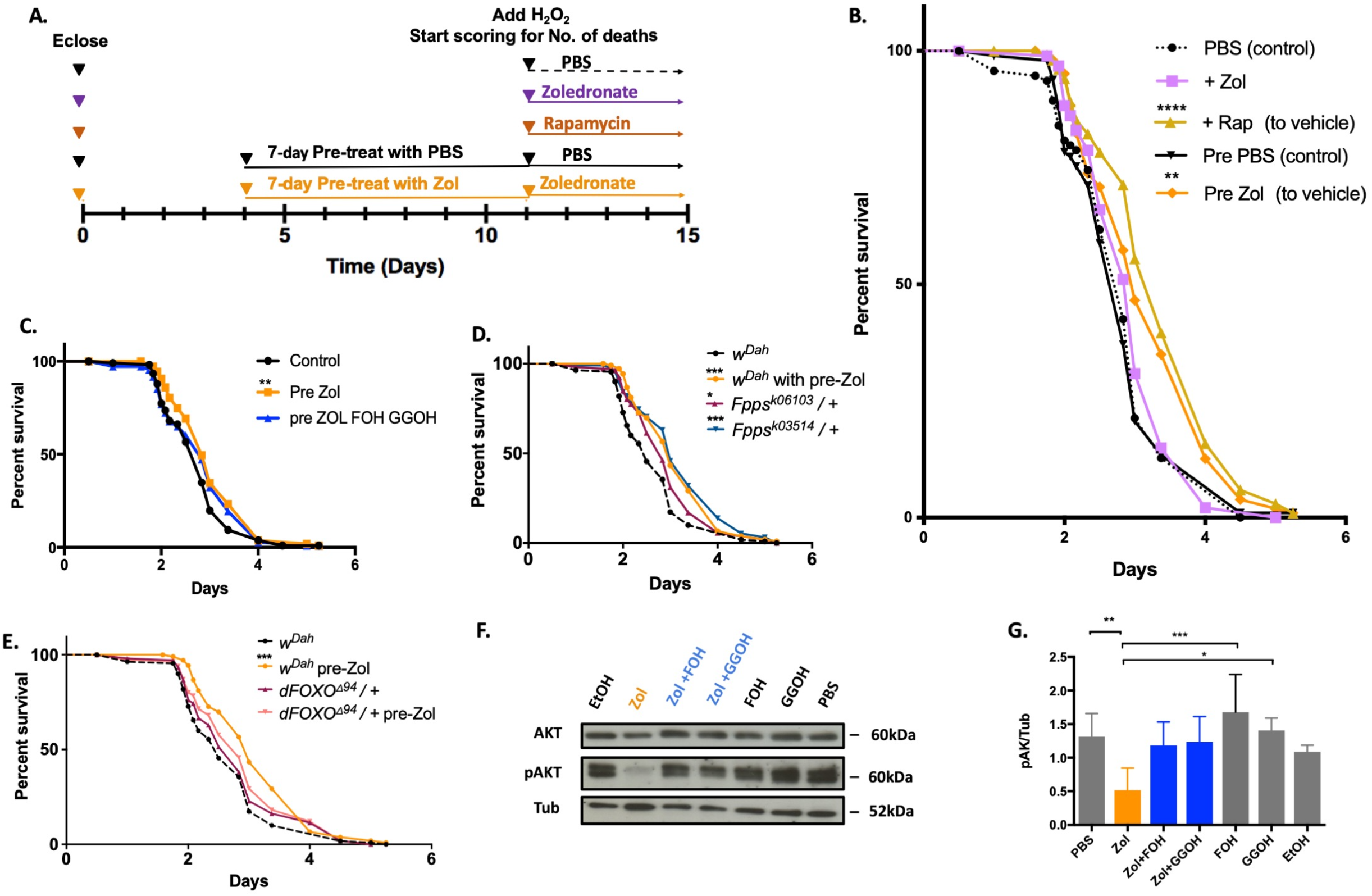
Zol increases lifespan of flies under oxidative stress through FOXO. (A) A schematic representation of the experimental design. (B) Survivorship of female *w*^*Dah*^ flies fed on food containing 5% H_2_O_2_. The flies were treated with 5% H_2_O_2_ in combination with PBS (vehicle) or 10μM Zol or 200μM rapamycin (Rap) both given at the same time than H_2_O_2_. Two groups of flies were pretreated for 7-days with 10μM Zol or PBS as vehicle before addition of H_2_O_2_ (C) Survivorship of female *w*^*Dah*^ flies fed with 5% H_2_O_2_. The flies were pre-treated PBS+EtOH (Vehicles), 10μM Zol, or 10μM Zol in combination with 330μM FOH & 330μM GGOH. (D) Survivorship of heterozygote female FPPS mutants: *Fpps*^*k06103*^/+ and *Fpps*^*k03514*^/+ fed with 5% H_2_O_2_. *w*^*Dah*^ flies either pre-treated PBS or 10μM Zol, were used as control (E) Survivorship of female dFOXO^Δ94/+^ flies treated with 5% H_2_O_2_. *w*^*Dah*^ flies either pre-treated with 10μM Zol or PBS (vehicle) were used as control. For all survival tests, 100 flies were used in each treatment group per test, experiments were repeated 3 times with different cohorts of flies. (F) A representative example of AKT and pAKT expression in whole flies fed with Zol (100μM) for 10 days, in presence or absence of FOH (330μM), GGOH (330μM). PBS and ethanol (EtOH) were used as vehicle control. (G) Quantification of expression level of pAKT normalised to tubulin in presence or absence of Zol (100μM), FOH (330μM) and GGOH (330μM) for 10 days analysed by ImageJ (n=3).

Taken together, these data suggest that treatment with Zol throughout adult life reduces intestinal dysplasia and permeability in *Drosophila*.

### Zol increases lifespan of flies exposed to oxidative stress through FOXO

To elucidate the mechanism of action mediating the beneficial effects of Zol and considering the role of GTPases in longevity we hypothesized that resistance to oxidative stress was partly responsible for the increased survivorship. Therefore, flies were challenged with food containing 5% H_2_O_2_, a treatment that reduces absolute lifespan and acts as a source of oxidative stress. Groups of adult, 11-day old female *w^Dah^* flies were subjected to 5% H_2_O_2_ at the same time as either Zol or Rapamycin used as a positive control (Emran, Yang, He, Zandveld, & Piper, 2014). We also tested another group of flies pre-treated with Zol for 7 days before exposure to 5% H_2_O_2_ (Fig. 5A). While the survival of flies treated with Rapamycin increased significantly compared to vehicle control, Zol-treatment did not improve survival when administered at the same time as the 5% H_2_O_2_ (Fig. 5B). By contrast, survival in response to H_2_O_2_ was significantly increased following pre-treatment with Zol for 7 days (Fig. 5B). To determine whether the increase was due to inhibition of FPPS, flies pre-treated with Zol were also treated with the downstream metabolites of the mevalonate pathway, Farnesyl farnesol (FOH) and Geranyl geranyol (GGOH), which bypass the block in FPPS inhibition. As expected, the survival advantage provided by Zol pre-treatment was abrogated in the presence of FOH and GGOH demonstrating that the extension in survival is due inhibition of the mevalonate pathway (Fig. 5C). To further strengthen the finding that inhibition of FPPS is responsible for the resistance to oxidative damage, *Fpps*^*k06103*^/+ and *Fpps*^*k03514*^/+ heterozygotes outcrossed into a *w*^*Dah*^ genetic background were also subjected to 5% H_2_O_2_ and their survival assessed. In these flies, survival was also significantly increased compared to *w*^*Dah*^ controls following exposure to H_2_O_2_-induced oxidative stress (Fig. 5D). Taken together these data suggest that the Zol confers resistance to oxidative stress and demonstrates that these effects are mediated by inhibition of the FPPS enzyme in the mevalonate pathway.

**Fig. 5.**
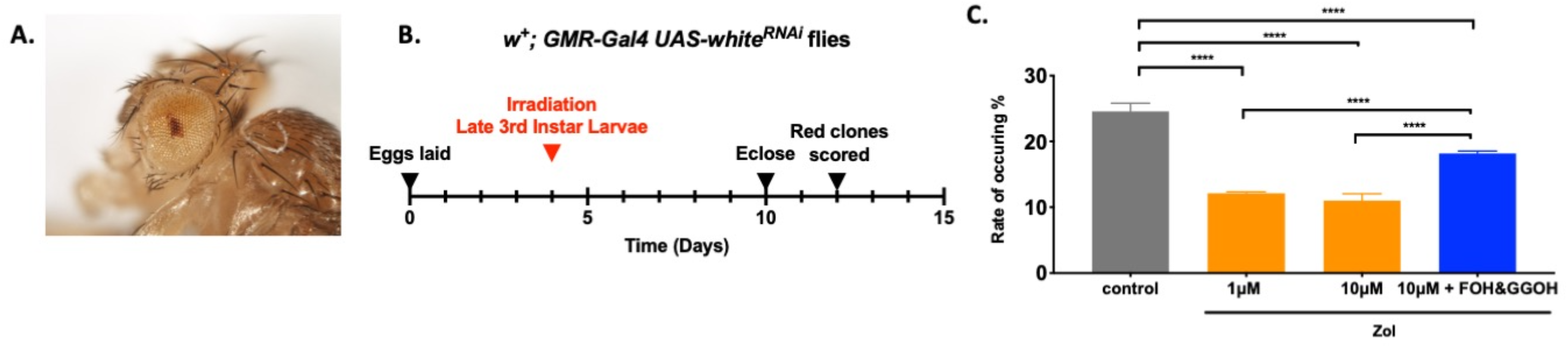
Zol enhances DNA damage repair in *Drosophila* upon irradiation. (A) Time line of experiment. (B) A representative image of an eye from a w+; GMR-Gal4, UAS-whiteRNAi fly containing a clone of cells in which DNA damage has led to the production of red pigment. (C) Percentage of eyes with red colonies upon 200Gy irradiation in homozygous *w*+; *GMR-Gal4,UAS-whiteRNAi* flies when treated with PBS (control), 1μM Zol, 10μM Zol, or 10μM Zol with 330μM FOH and GGOH. Data analysed by one way-ANOVA and Boneferroni post-hoc test. ***p<0.001, ****p<0.0001

To determine the mechanism of action downstream of the mevalonate pathway, we hypothesized that Zol may be affecting the activity of dFOXO, a factor regulating oxidative stress signals. To determine whether Zol protects *Drosophila* from oxidative damage through dFOXO, heterozygous *dFOXO*^*Δ94*^/+ loss of function mutations (Slack, Giannakou, Foley, Goss, & Partridge, 2011) and *w*^*Dah*^ controls were pre-treated with either Zol or vehicle control-containing food for 7 days before exposure to food containing 5% H_2_O_2_. While *w*^*Dah*^ pre-treated with Zol showed a significant increase in lifespan, *dFOXO* ^Δ*94/+*^ flies treated in the same way did not show any beneficial effect (Fig. 5E), suggesting dFOXO is required for Zol action on survival.

Given that pAKT is a known regulator of FOXO (Accili & Arden, 2004; Das, Suman, Alatassi, Ankem, & Damodaran, 2016) we next determined levels of pAKT following treatment with Zol and observed a significant decrease in pAKT levels (Fig. 4F and S2). Furthermore, these effects were reversed by the addition of FOH and GGOH (Fig. 5 F-G), suggesting that inhibition of pAKT is dependent on the mevalonate pathway.

Taken together these data suggest that Zol confers resistance to oxidative damage via inhibition of the mevalonate pathway through a mechanism that requires dFOXO for its activity.

### Zol enhances DNA damage repair in *Drosophila* upon irradiation

DNA damage and the resulting mutations induced by ionizing radiation are at least partly caused by increased oxidative stress (Azzam, Jay-Gerin, & Pain, 2012; Riley, 1994). We therefore wanted to determine whether the ability of Zol to protect against H_2_O_2_-induced oxidative stress would also reduce the frequency of DNA damage induced by X-ray irradiation. In order to visualise DNA damage in an *in vivo* environment, we developed an assay in which endogenously expressed mRNA from the wild type *white* locus is knocked down by an *in vivo* RNAi hairpin-loop expressed within the cells of the future eye. Where knock down is successful, *white* mRNA is destroyed and very little pigment is produced, resulting in pale yellow eye pigmentation. Where the DNA encoding components of the *GMR-Gal4,UAS-white*^*RNAi*^ expression system are mutated, *white* mRNA is not destroyed and wild type levels of red pigment are produced to give a readily recognisable red eye clone (Fig 6A) in adult *Drosophila*. In order to assess DNA damage levels using this reporter, larvae heterozygous for the *GMR-Gal4,UAS-white*^*RNAi*^ reporter were raised on food containing either carrier controls, 1μM Zol, 10μM Zol or a combination of10μM Zol and 330μM FOH and 330μM GGOH. Larvae were irradiated 96 hours after hatching using one dose of 200Gy of X-ray irradiation (Fig. 6C). Strikingly, *GMR-Gal4,UAS-white*^*RNAi*^/+ flies contained significantly lower frequency of red-marked mutated cells when grown on 1 or 10μM Zol-containing food compared to controls. However, when *Drosophila* were treated with 10μM Zol in combination with FOH and GGOH, the reduction in DNA damage achieved by Zol was abrogated and the frequency of red-marked mutant cells clones detected was no longer significantly different to the controls (Fig 6D). These data suggest that Zol is also able to protect individuals from the accumulation of mutations, via mechanism(s) that depend on the mevalonate pathway.

## DISCUSSION

In this study we show that Zol has properties of a geroprotector, an activity mediated by its inhibition of FPPS. We show that Zol extends the lifespan and healthspan of *Drosophila* in absence of mineralised bone-like structures and demonstrate that it confers resistance to oxidative damage via the inhibition of FPPS in the mevalonate pathway. Zol requires dFOXO for its action with this factor being modulated via changes in the levels of pAkt. These *in vivo* findings are in line with our previous work in human mesenchymal stem cells (hMSC) where we have shown that Zol was able to reduce the accumulation of DNA damage caused by cellular ageing or irradiation via a mechanism requiring FOXO3a, as well as reducing the accumulation of senescent markers p16 and p21 (Misra et al., 2015). FOXO is a key player in ageing and has been shown to regulate several of its hallmarks including DNA damage, senescence and changes in mitochondrial function (Borch Jensen, Qi, Riley, Rabkina, & Jasper, 2017; de Keizer et al., 2010; Qi et al., 2015; Tran et al., 2002; Tsai, Chung, Takahashi, Xu, & Hu, 2008). Whilst *Drosophila* have a single FOXO gene (dFOXO), the human genome is more complex and encodes four FOXO different proteins, with polymorphisms in FOXO3a having been associated with exceptional longevity (Donlon et al., 2012; J. C. Lee et al., 2013). In addition, FOXO3a activity has been shown to reduce the effects of reactive oxygen species (ROS) production in multiple ways. For example, its expression improves the fidelity of DNA damage repair by arresting the cell cycle to allow the repair of damaged DNA (Fluteau et al., 2015; Gurkar et al., 2018; Tran et al., 2002; Tsai et al., 2008). In addition FOXO3a activation results in the repression of a large number of nuclear-encoded genes with mitochondrial function (Ferber et al., 2012), (Ferber et al., 2012). As most intrinsic ROS are produced by the respiratory complexes located in the inner mitochondrial membrane, these changes in mitochondrial activity may directly influence the levels of ROS production *in vivo*. More work is required to understand in detail which of these mechanisms and molecular pathways Zol modulates via FOXO. A detailed analysis of each individual tissue in a mammalian system is also required as there are important differences in response to oxidative stress not only among tissues but even within regions of the same tissues (Stefanatos & Sanz, 2018).

One observation we made is that the effects of Zol varied depending on time of administration, dose and sex differences that may be partly due to alternate drug uptake by males and females. In females, egg production requires higher levels of nutritional input than required by males, resulting in increased food consumption and, presumably, increased drug uptake (Camus, Huang, Reuter, & Fowler, 2018). By contrast, in the FPPS mutants, where the action of the enzyme is disrupted independently of drug uptake, similar lifespan extension is observed suggesting that the effect of mevalonate pathway inhibition on lifespan is unlikely to be sexually dimorphic.

In addition to differences in lifespan extension, it is intriguing to note that indicators of improved health such as climbing ability and intestinal dysplasia occur even at high doses of Zol that reduce absolute lifespan (compare Figure 1B 10μM with Figure 3A, 63day 10μM), demonstrating that drug treatment is able to increase healthspan independently of absolute lifespan. This observation is in line with the finding of others which shows that healthspan and lifespan are not necessarily related (Fischer et al., 2016). For example, Nicotinamide has recently been shown to improve aspects of healthspan but not lifespan (Mitchell et al., 2018). This disconnect between healthspan and lifespan may indicate that organisms can cope with the accumulation of a certain number of defects which are sufficient to negatively affect healthspan but which do not lead directly to death. Being able to reduce these healthspan-associated deficits is of particular interest from a translational perspective as they contribute to morbidity and poor health and are responsible for significant healthcare costs.

Another notable aspect of Zol treatment in *Drosophila* is the improvement in lifespan and healthspan measures following treatment that begins only in middle age (Figure 1E-F). This treatment profile mirrors that of women affected by osteoporosis who also generally begin Zol treatment post-menopausally. In both *Drosophila* and humans the effect of Zol on survival is only detectable sometime after initial treatment (Lyles et al., 2007). On average it takes 16 months before an improvement in survival is observed in postmenopausal women (Lyles et al., 2007) and approximately 14 days in *Drosophila* (Figure 1F). The reasons for this delayed phenotypic response are unclear but may reflect an ability by Zol to improve biological processes only when they are mildly dysregulated. However, when those mechanisms are either working at healthy levels or when their dysregulation exceeds a threshold then Zol treatment is either not needed or no longer sufficient to maintain function. Consistent with this model, Zol did not have any effect on climbing activity in geriatric flies at 70 days of age – when no activity was detected in any of the populations tested. It is possible that a similar process may also explain why extension of lifespan was not observed in *Zmpste24*^−/−^ mice. In this model of Hutchinson-Guilford progeria syndrome Zol alone does not modify premature ageing while combined treatment with both Zol and statins (which inhibit the same pathway) is able to extend lifespan (Varela et al., 2008).

While the effects of Zol on lifespan and healthspan in humans may appear to be limited, it should be noted that these effects are observed following treatment with just a single yearly dose. Retrospective analysis of patients taking Zol has not only shown a marked increase in survival but also a reduction in the frequency of pneumonia and cardiovascular events, suggesting broader effects unrelated to the musculoskeletal system (Colón-Emeric et al., 2010). In addition, patients treated with Zol and admitted to hospital for intensive / critical care for a condition not related to osteoporosis showed reduced mortality rates relative to controls (5.2% vs 9.1% respectively). Furthermore, the patients previously treated with Zol required 30% shorter in-patient care despite being older than the control group and having higher comorbidity index (P. Lee et al., 2016a).

Older people with frailty and multimorbidity have a reduced ability to respond to adverse events and often lose independence following major health-related incidents. Such events have significant consequences for follow up health and social care costs (Liotta et al., 2019; National Audit Office, 2016). Indeed, there has been an increase of 18% in the number of emergency admissions of older patients between 2010/11 and 2014/15 in the UK with those patients over the age of 65 now accounting for 62% of total bed days spent in hospital. Considering average cost for a patient to stay in an NHS ward is up to £400 per day, the financial and societal benefits of improving resilience in older people and reducing length of hospital stay are huge (NAO, 2016).

In conclusion we have shown that inhibition of FPPS by Zol modulates mechanisms of ageing to extend lifespan and healthspan *in vivo* – an effect that is independent of its effects on bone. These findings are in line with the unexplained improved survival rates that have been reported recently for patients being treated with Zol – findings that highlight the substantial benefit Zol can potentially provide with only a single yearly treatment. Studies in mammalian models are now required to understand the effects of Zol on specific tissues, to define which of the FOXO-regulated mechanisms are at play and whether an infrequent administration of Zol is sufficient to elicit the beneficial effects. Given that Zol is off-patent, available at low cost and displays a well understood safety profile featuring minimal side effects, we suggest that repurposing studies seeking to widen the use of Zol are likely to identify great potential for the improvement of healthspan and resilience in older people.

## EXPERIMENTAL METHODS

### Fly Stocks and Husbandry

The *white*^Dahomey^ (*w*^*Dah*^) stock has been maintained in large population cages with overlapping generations since 1970 (Grandison, Wong, Bass, Partridge, & Piper, 2009) and was a kind gift of the Partridge lab. *y*^*1*^ *w*^*67*^*c*^*23*^; *P*{*w*^+*mC*^=*lacW}Fppsk*^*06103*^/*CyO* and *y*^*1*^*w*^*67*^*c*^*23*^; *raw*^*k03514*^*P{w*^+*mC*^=*lacW}Fpps*^*k03514*^/*CyO* (Spradling et al., 1999) and *w*^*1118*^; *foxo*^*Δ94*^/*TM6B*, *Tb*^*1*^ (C Slack, Giannakou, Foley, Goss, & Partridge, 2011) were obtained from the Bloomington *Drosophila* Stock Centre. *w; Su(H)-lacZ, esg-Gal4,UAS-GFP/CyO* expresses GFP in intestinal stem cells and progenitor cells and was a kind gift of Leanne Jones (Choi, Kim, Yang, Kim, & Yoo, 2008). *w+; GMR-Gal4,UAS-white^RNAi^* is a recombinant between *GMR-Gal4* (Hui Liu, Ma, & Moses, 1996) and GD30033, an *in vivo* hairpin loop RNAi construct targeting *white* mRNA from the Vienna *Drosophila* Resource Center (Dietzl et al., 2007).

All flies were maintained in a 12:12h light-dark cycle on standard yeast, molasses, cornmeal, agar food at 18°C or 25°C. Drug-containing food was prepared using freshly cooked molten fly food cooled to 60°C. Zol (kindly provided by Hal Hebetino, University of Rochester), GGOH (Sigma Aldrich, Dorset, U.K.) and FOH (Sigma Aldrich, Dorset, U.K.) or carrier(s) were added, mixed and immediately poured into standard vials and bottles so as to minimize exposure of drug to high temperatures. The continued activity of Zol following brief exposure to this temperature was confirmed molecularly (data not shown). Drug food was stored at 4°C until use and unused food discarded after 3 weeks.

For lifespan experiments overnight embryo collections from approximately 200 adult flies were collected on apple juice agar plates, washed with water and 32μl of embryos transferred into food bottles. Flies that subsequently ecclosed within the first 24hrs of the first to emerge were discarded with those subsequently ecclosing over an 16-20 hour overnight collection window being used for experiments. These later flies were transferred into new food bottles without the use of CO_2_ and incubated for 3-4 days. Flies were then sorted by gender and counted while minimizing CO_2_ exposure.

### Longevity Assay and H_2_O_2_ Survival Assays

For longevity assays, 20 gender matched flies were maintained in vials of food and transferred into new vials every 2-3 days. At every transfer numbers of dead or censored flies were recorded. Unless specifically specified 100 flies were analysed for each experimental with three independent experimental replicates. To test survival in presence of hydrogen peroxide, adult flies of the appropriate genotype and pre-treatment were first starved in vials containing 1% agar for 3 hours and then transferred to standard food containing 5% H_2_O_2_ and the appropriate drugs / vehicle. Death was then scored at 12, 24 and 36 hours and then every 2 hours for the next 48 hours. If 100% mortality was not reached by that point, scoring was continued every 12 hours.

### Healthspan assays

For rapid iterative negative geotaxis (RING), assays flies we kept in groups of ~150. The day before an experiment, flies were sorted into groups of 20 and transferred into separate vials following brief CO_2_ anaesthesia. Flies were then transferred into 25ml strippette (Fisher Scientific) and after 1 minute, the strippette was tapped to knock flies to the bottom and initiate the negative geotaxic response. After 10 seconds a photograph was taken and the position of each fly, and hence the distance climbed, was recorded. Twenty flies were assessed from each condition with each experiment repeated 3 times. Flies that climbed above 10cm were classified as ‘high climbers’.

For gut barrier integrity assays, and food uptake assays *Drosophila* food containing 2.5% erioglaucine disodium salt (Sigma Aldrich) and drugs (as appropriate) was fed to flies for 9 hours. For gut integrity assays flies were scored for uniform blue coloration beyond the GI tract (‘Smurf-ness’). For food uptake assays 60 flies were homogenised in 1ml water, passed through a 0.2μm filter and measured for OD629 using a Nanodrop Spectrophotometer ND-1000 (Labtech International, Sussex, UK) previously blanked with flies fed with standard *Drosophila* food.

### DNA damage assay

An overnight egg collection of *w*+, *GMR-Gal4 UAS-white*^*RNAi*^ adults outcrossed to OreR were allowed to develop for 96 hours at 25°C. These heterozygous third instar larvae were then irradiated with 200Gy using a Torrex Cabinet X-ray system (Faxitron X-ray, Arizona, USA) and adult flies that subsequently ecclosed were scored for the frequency of red *w*^+^ clones.

### *Drosophila* midgut analysis

*Midguts from Su(H)-lacZ; esg-Gal4,UAS-GFP/CyO* adults were dissected in cold PBS and fixed in PBS+4% formaldehyde and blocked in PBST (PBS, 0.1% Triton X-100 and 1% BSA). Antibodies included Chicken anti-GFP 1:4000 (Abcam, Cambridge, UK) and anti-Phospho-histone H3 1:100 (New England Biolabs, Ipswich, USA) and secondary antibodies AlexaFluor^TM^ 488 goat anti-chicken and AlexaFluor^TM^ 594 goat anti-rabbit IgG (Life Technologies, Oregon, USA) were used. Stained guts were mounted with Fluoroshield^TM^ with DAPI (Sigma Aldrich) and Grace Bio-Labs SecureSeal™ imaging spacers. Samples were imaged using Perkin Elmer Spinning Disk confocal microscope with a 40X objective. Images were then processed using Image J.

### Protein extraction and western blot

30 flies of the appropriate genotype were snap frozen in liquid N_2_ and crushed using an Eppendorf pestle in 200μl ice-old lysis buffer (50mM Tris HCl pH 7.4, 250mM NaCl, 5mM EDTA and 0.003% Triton X-100) with protease inhibitor (cOmplete Mini, EDTA-free, ROCHE, Sussex, UK). After 30mins incubation at 4°C extracts were centrifuged at 13,000rpm, 4°C for 30mins and supernatants stored at −20°C.

40-60μg of protein was electrophoresed through 4–15% Mini-PROTEAN® TGX™ Precast Protein Gels (Bio-Rad, Hertfordshire, UK) and transferred onto polyvinylidene difluoride (PVDF) membrane (Immobilon-P Membrane, Millipore, Hertfordshire, UK) using standard techniques. Primary antibodies used were anti-pAKT 1:500, anti-AKT 1:500, anti-tubulin 1:1000 (Cell Signaling Technology, Denver, Massachusetts). HRP conjugated secondary antibodies were used at 1:10,000 and visualised using ECL (GE Healthcare, Buckinghamshire, UK), high performance chemiluminescence film (Amersham Hyperfilm ECL, GE Healthcare) and an Optimax 2010 X-ray film processor (PROTEC, Oberstenfeld, Germany).

### Statistical Analysis

Statistical analysis was performed on GraphPad Prism 7. Paired data was analysed using non-parametric unpaired Student’s t-test unless otherwise stated. For multiple comparisons, ordinary one-way analysis of variance (ANOVA) and two-way ANOVA was used followed by Bonferroni’s Sidak’s multiple comparison post-hoc tests. Survival data was analysed using Log-rank (Mantel-Cox) test. The death curve generated was then analysed using the Gehan-Breslow-Wilcoxon test to identify significant differences in organism survival between treatment groups. All data is expressed as mean ± standard deviation (SD). A difference was stated to be statistically significant if the p value was <0.05 (*p<0.05; **p<0.01; ***p<0.001; ****p<0.0001).

## Acknowledgements

We would like to thank Leanne Joes for fly stocks and valuable advice, Kath Whitley and fly room technical support staff for assistance with fly food preparation. We would also like to thank the Bloomington and Vienna *Drosophila* stock centres for fly stocks and Hal Hebetino and Graham Russell for donating zoledronate and for helpful discussions.

## Author Contributions

IB, MZ conceived the idea, analysed the data, wrote the manuscript ZC performed the experiments, analysed the data and wrote the manuscript. All authors designed the experiments, reviewed and approved the manuscript.

## Data Availability

The data that support the findings of this study are available from the corresponding author upon reasonable request.

