## Supplementary figure 1 & 2 for "Zoledronate extends healthspan and survival via the mevalonate pathway in a FOXO-dependent manner"

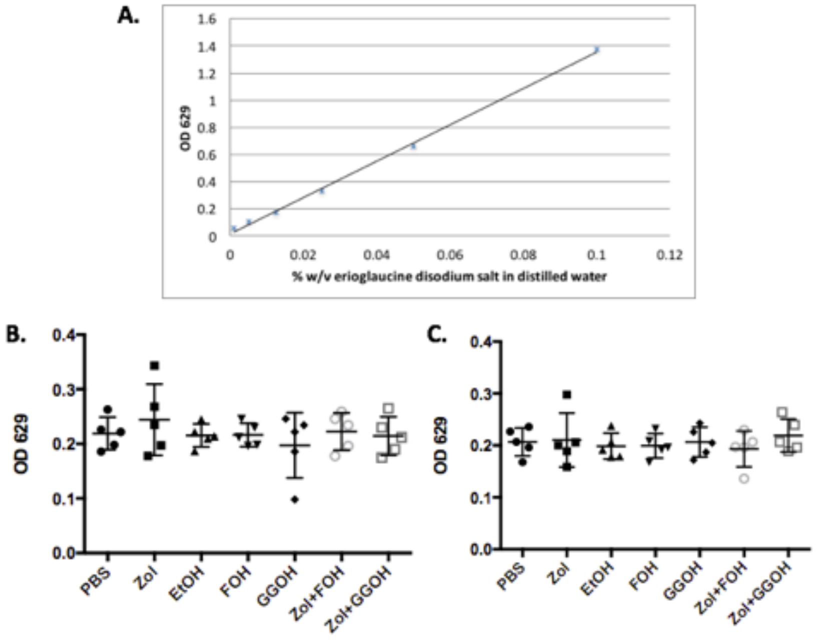


Fig. S1 **<u>Food intake level was not affect with addition of Zol, FOH or GGOH in their diet</u>**

*(A) Standard curve between concentration of erioglaucine disodium salt in distilled water to OD 629 reading. (B) Quantification of blue colouring in Drosophila when flies were fed on 'blue food' containing different molecules for half an hour (n=5). (C) Quantification of blue colouring in Drosophila when flies were subjected to food containing different molecule combinations for 7 days prior to start of the assay (n=5). Data are expressed as mean ±SD and were analysed by one way ANOVA and Bonferroni post-test for multiple comparisons *p<0.05, **p<0.01, ***p<0.001, ****p<0.0001.*

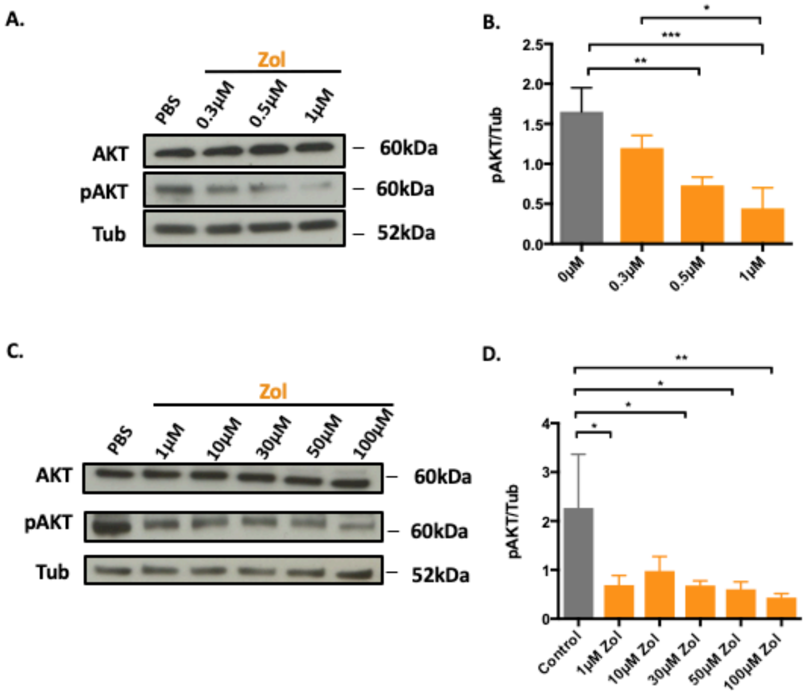


Fig. S2 **Zol reduces pAKT level in *Drosophila* in a dose-dependant way**

*(A) A representative example of AKT and pAKT expression in whole flies fed on food containing 0.3-1µM Zol for 10 days. Group of flies were also fed with food containing PBS only for control. (B) Quantification of expression level of pAKT normalised to tubulin (Tub) in presence or absence of Zol (0.3-1µM) in Drosophila food for 10 days analysed by imageJ (n=3). (C) A representative example of AKT and pAKT expression in whole flies fed on food containing 1-100µM Zol for 10 days. Group of flies were also fed with food containing PBS only for control. (D) Quantification of expression level of pAKT normalised to tubulin (Tub) in presence or absence of Zol (1-100µM) in Drosophila food for 10 days analysed by imageJ (n=3).*
